# Impact of native-like lipid membranes on the architecture and contractility of actomyosin networks

**DOI:** 10.1101/2022.07.22.501094

**Authors:** Nils L. Liebe, Ingo Mey, Loan Vuong, Burkhard Geil, Andreas Janshoff, Claudia Steinem

## Abstract

The connection between the actomyosin cortex and the plasma membrane of eukaryotic cells is investigated by creating a versatile, near-native model system that allows studying the architecture and contractility of the cortex as a function of lipid composition. We found that the concentration of phosphatidylserine, a characteristic lipid of the inner leaflet of mammalian plasma membranes, plays a pivotal role in the binding of the membrane-cytoskeleton linker protein ezrin and the resulting contractile behavior of an adjacent actin network. In addition to the specific receptor lipid for ezrin, i.e., PtdIns[4,5]P_2_ cross-linking the network to the inner leaflet, the presence of phosphatidylserine in the membrane is critical to enhancing the binding of ezrin to PtdIns[4,5]P_2_ and allows rapid local actin contraction at physiologically relevant concentrations in the regime of 1-3 mol% PtdIns[4,5]P_2_. In the presence of phosphatidylserine, the additional negative charges in the membrane may induce enhanced sliding of the filaments on the membrane surface due to repulsive interactions between F-actin and the bilayer readily leading to the emergence of contraction foci. Conversely, if phosphatidylserine is replaced by an increased PtdIns[4,5]P_2_ concentration of 5 or 8 mol%, a highly connected but non-contracting actin network is observed.

---

Cell shape and cellular dynamics predominately rely on the active mechanical properties of the actomyosin cortex. The cortex of nonmuscle cells, which is about 200 nm thick, is formed by an apolar, disordered network of transiently crosslinked actin filaments pre-stressed by the presence of nonprocessive myosin II motors and attached to the plasma membrane by specific anchors (1, 2). This pre-stress gives rise to a substantial tension that is essential for changing the cell’s shape during migration, maturation and division. Cells regulate contractility of the actomyosin cortex locally and transiently by altering the density of cross-links and motor activity. These two degrees of freedom experience a boundary, the cell membrane. Tethering the actin mesh to the plasma membrane is realized by specific proteins such as those of the ezrin/radixin/moesin (ERM) protein family (3), which are evolutionary highly conserved and tissue specific. Among them, ezrin is found in epithelial cells (4), where it links the plasma membrane via PtdIns[4,5]P_2_ to its N-terminal domain (N-ERMAD) (5) and F-actin to the C-terminal domain (CERMAD), respectively (4, 6). Ezrin activation from a cytosolic dormant conformation to an activated state occurs upon first binding to PtdIns[4,5]P_2_ followed by phosphorylation of threonine-567 (7, 8). In the active conformation, it serves as a passive, reversible linker between F-actin and the plasma membrane, whereas myosin II motors actively reorganize the network and thereby generat tension in the cortex (9, 10). This cortical tension depends on the local F-actin organization (2) as well as the membrane-cortex linkage (11).

To entangle the different contributions of membrane, linker proteins and myosin motors on the actomyosin organization and contractility, minimal actin cortices (MACs) have been developed (12). Besides three-dimensional model systems based on giant unilamellar vesicles (GUVs) with actin cortices attached either to the outside (13–15) or the inside of the GUV (15–18), planar supported lipid bilayers (SLBs) turned out to be particularly suitable as they allow for highresolution fluorescence microscopy analysis of actin network organization and dynamics in the context of a two dimensional natural boundary.

Several strategies were followed to link F-actin to SLBs (11, 19–24). However, even though these artificial planar membrane systems provide valuable information about the architecture and contractility of membrane-attached actomyosin cortices, none of them reflect the lipid composition and membrane-anchoring found at the plasma membrane of epithelial cells. For instance, the lipid composition of the plasma membrane is highly asymmetric and this uneven distribution of lipids is actively maintained (25–27). As a result, the extracellular leaflet is composed mostly of phosphatidylcholine and sphingomyelin, while only the cytoplasmic leaflet contains phosphatidylserine. The influence of the associated negative charge density provided by phosphatidylserine on the architecture and dynamics of the actomyosin cortex has been as yet fully neglected.

Here, we show that a lipid composition resembling the inner leaflet of a eukaryotic mammalian cell harboring in particular negatively charged phosphatidylserine (PS) and phosphoinositolphosphate (PtdIns[4,5]P_2_) is essential for proper function of the cortex, i.e., connectivity paired with contractility. Therefore, we formed MACs in which both the native linkage to the membrane as well as the native lipid composition of the inner leaflet of the plasma membrane of epithelial cells are reproduced. We found that the F-actin architecture and contraction dynamics strongly depends on the presence of PtdIns[4,5]P_2_ and PS. The emergence of large scale structures in membrane-associated actin networks depends on the PtdIns[4,5]P_2_ and PS content. Only in the presence of PS, the connectivity and contractility is rapid, but a global collapse of the network is avoided. This finding has potential consequences for the function of the cortex as a pre-stressed network capable of maintaining mechanical homeostasis but also dynamically controlling the shape of cells in situations that require a rapid response such as adhesion, migration, division and tissue formation.

## Results

### The impact of POPS and PtdIns[4,5]P_2_ on the architecture of the minimal actin cortex

Minimal actin cortices (MACs) were produced step-by step on top of a supported lipid bilayer (SLB) (Fig. 1A) (29, 30). Planar SLBs composed of either POPC and PtdIns[4,5]P_2_ or POPC, PtdIns[4,5]P_2_ and POPS were prepared on either glass or silicon substrates and imaged by means of fluorescence microscopy (Fig. 1B). Afterwards, the active mutant of ezrin (ezrin T567D) was added, which specifically binds to PtdIns[4,5]P_2_ (Fig. 1C, Fig. S1). Ezrin forms reversible cross-links between the actin filaments and the membrane (6, 28, 29). Confocal laser scanning micrographs taken after addition of pre-polymerized F-actin reveal quasi two-dimensional networks attached to the membrane (Fig. 1D). However, at 2 mol% of PtdIns[4,5]P_2_, the F-actin network appears to be much denser in the presence of POPS than in its absence (Fig. 1D).

**Fig. 1.**
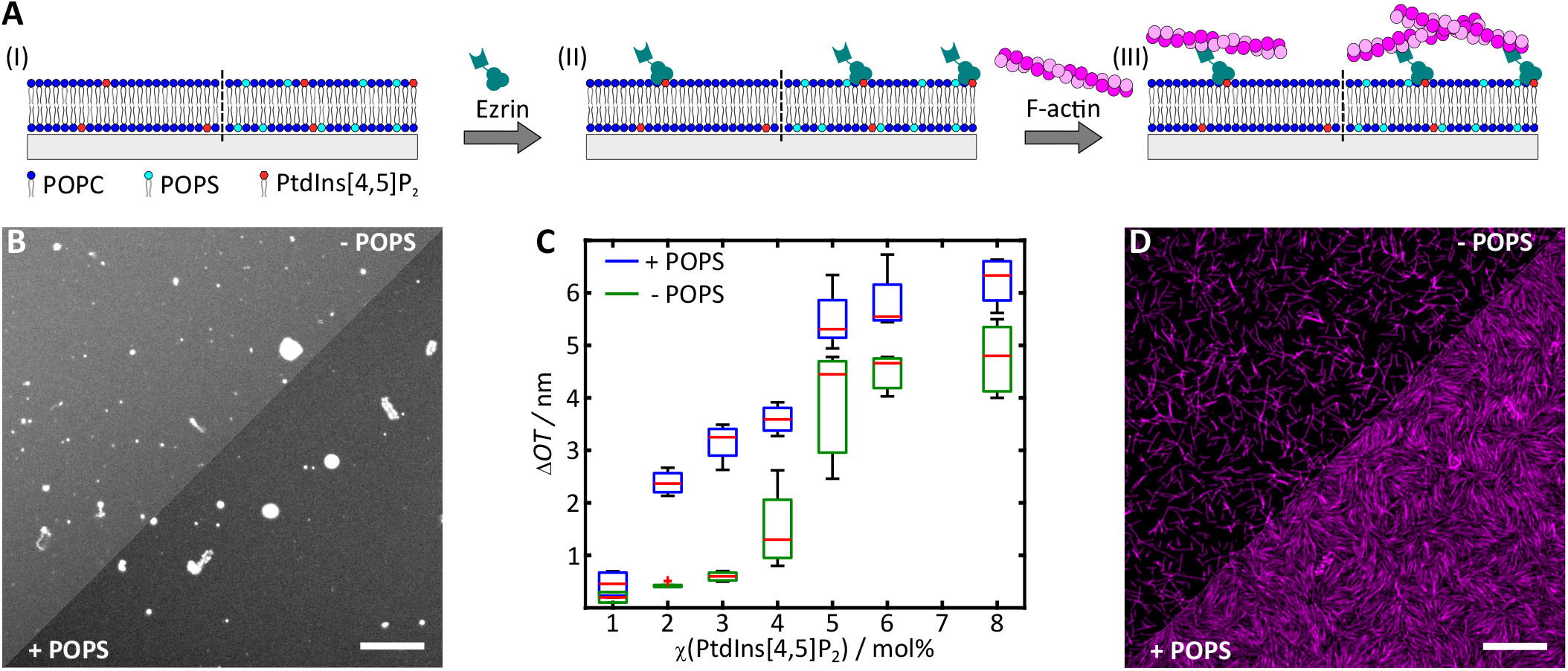
Binding of minimal actin cortices (MACs) to solid supported lipid bilayers (SLBs) doped with and without POPS. (**A**) Schematic illustration of the key steps of MAC preparation. (I) SLB formation by spreading SUVs composed of POPC/PtdIns[4,5]P_2_ (left) or POPC/PtdIns[4,5]P_2_/POPS (right) on hydrophilic surfaces. (II) Specific binding of the F-actin-membrane linker ezrin (active mutant T567D) to the receptor lipid PtdIns[4,5]P_2_ and (III) coupling of pre-polymerized F-actin filaments. (**B**) Fluorescence micrographs of a glass supported lipid bilayer composed of POPC/PtdIns[4,5]P_2_/ATTO 390-DPPE (upper part, 97.6:2:0.4) and POPC/PtdIns[4,5]P_2_/POPS/ATTO 390-DPPE (lower part, 80.6:2:17:0.4) (gray). (**C**) Change in optical thickness (Δ*OT*) caused by ezrin T567D binding to SLBs without (green) and doped with 17 mol% POPS (blue) as a function of the PtdIns[4,5]P_2_ content. For the analysis *m* experiments were performed. 1 mol% (*m* = 4, *m* = 2), 2 mol% (*m* = 4, *m* = 5), 3 mol% (*m* = 4, *m* = 5), 4 mol% (*m* = 4, *m* = 4), 5 mol% (*m* = 6, *m* = 3), 6 mol% (*m* = 4, *m* = 3) and 8 mol% (*m* = 4, *m* = 7). Δ*OT* data in the absence of POPS were taken from Nöding et al. (28). Boxes range from 25^th^ to 75^th^ percentiles of the sample, while whiskers represent the most extreme data points not considered as outliers. Medians are shown as red horizontals within the boxes. (**D**) Fluorescence micrographs of pre-polymerized F-actin bound to an SLB composed of POPC/PtdIns[4,5]P_2_/ATTO 390-DPPE (upper part, 97.6:2:0.4) and POPC/ PtdIns[4,5]P_2_/POPS/ATTO 390-DPPE (lower part, 80.6:2:17:0.4) after incubation with ezrin T567D. Scale bars: 10 µm (B, D).

Besides a denser network, F-actin bound to POPS-containing membranes tends to form local domains of nematic alignment (Fig. 1D). This finding might be indicative of repulsive interactions among the actin filaments and between the filaments and the negative charges provided by POPS fostering filament alignment and being responsible for the emergence of large scale patterns. The thermal motion of the filaments on the surface leads to steric repulsion as soon as two filaments meet in an angle, and consequently to a parallel/antiparallel alignment that results in the observed nematic ordering. In order to quantify the increase in local order with the surface concentration of actin, we used an approach described by Seara et al. (31) and determined the coarse-grained nematic order parameter *q* = 2(cos^2^*θ*−1*/*2) (Fig. 2A1-3). Seara et al. (31) used a crowding agent to accumulate F-actin on a membrane surface and found that the network continuously changes from an isotropic to a nematic phase as the F-actin concentration on the surface increases. To relate the average nematic order parameter to the network density, we applied a “*tube filter*” analysis to the fluorescence micrographs (Fig. 2B1) to skeletonize the network (Fig. 2B2). From the skeletonized images we were then able to determine the skeleton network density, defined as the ratio of filamentous pixels to all image pixels and the filament intersections reported as node density (Fig. 2B3, red crosses) (28).

**Fig. 2.**
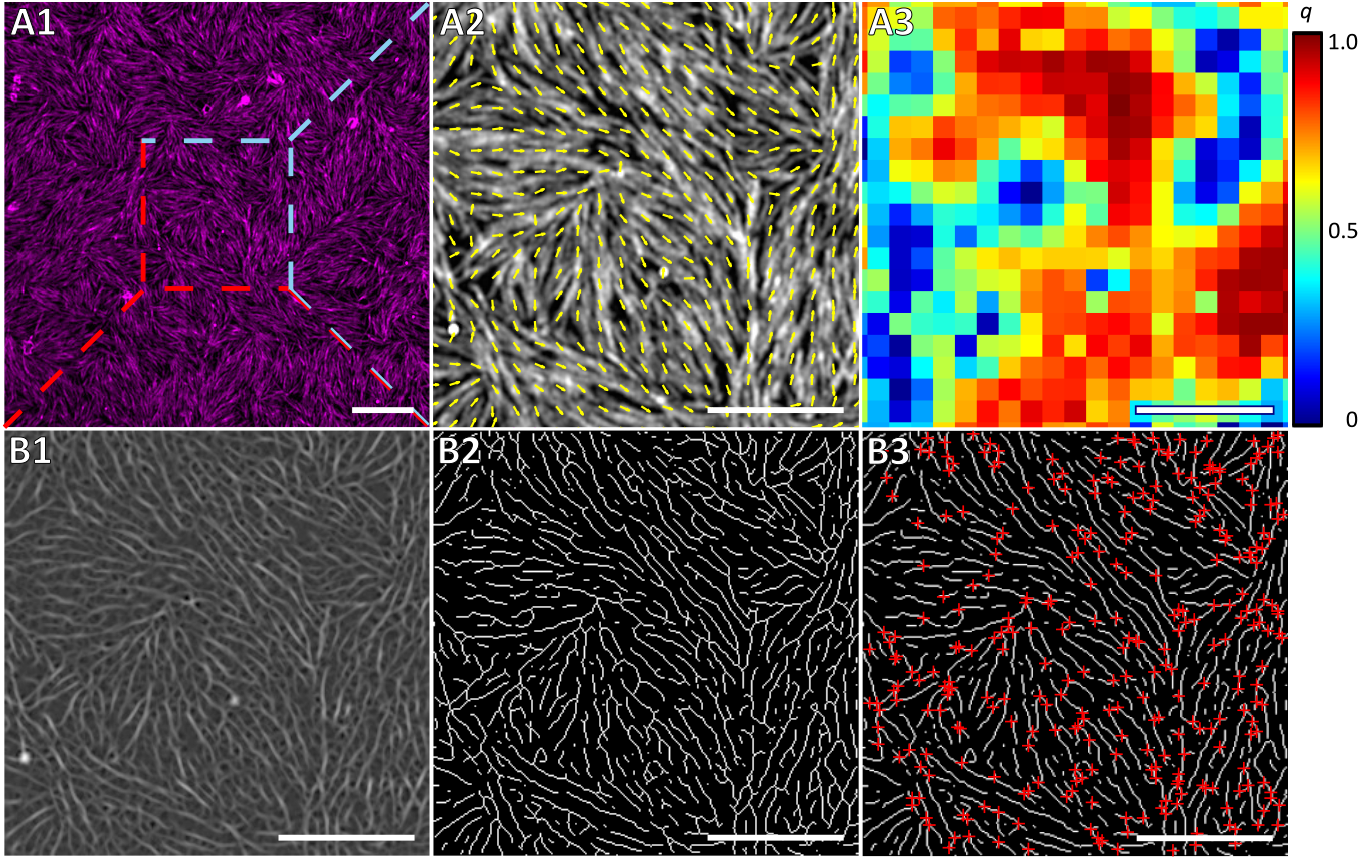
Self-organization of membrane bound minimal actin cortices. (**A1**) Fluorescence micrograph of an F-actin network (magenta) bound to a POPS doped SLB showing a nematic self-organization. (**A2**) Overlay of the alignment vector field (yellow arrows) and the fluorescence image (A1, shown in grey) representing the nematic phase direction within the membrane bound F-actin network. (**A3**) Heat map of the scalar nematic order parameter (*q*) for the F-actin network shown in A1. (**B1**) “*Tube filter* “analysis and resulting skeletonization (**B2**) of a zoom in of A1 (marked with a dashed white square). (**B3**) Overlay of the skeletonized F-actin network with detected nodes represented as red crosses. Membrane composition: POPC/PtdIns[4,5]P_2_/POPS/ATTO 390-DPPE (80.6:2:17:0.4); scale bars: 10 µm (A1), 5 µm (A2-B3).

The network density was controlled by the surface concentration of PtdIns[4,5]P_2_ limiting the amount of available ezrin cross-linkers for the actin network. We varied the PtdIns[4,5]P_2_ concentration within the physiological relevant concentration range of 1-3 mol% PtdIns[4,5]P_2_. Together with 17 mol% POPS, this lipid mixture mirrors the essential composition of the inner human plasma membrane containing PtdIns[4,5]P_2_ (1-5 mol%) and POPS (12-20 mol%) (26, 32, 33). In the absence of POPS and at lowest PtdIns[4,5]P_2_ concentration, only few filaments bind to the ezrin-decorated membrane (Fig. 3A, top left). With increasing PtdIns[4,5]P_2_ concentration, the surface concentration of actin filaments increases significantly (Fig. 3A, top middle and right). Naturally, the increased network density is also reflected in the skeleton network density (Fig. 3D). In the presence of POPS, the overall filament density considerably increases for 1 and 2 mol% PtdIns[4,5]P_2_. Already at 1 mol% PtdIns[4,5]P_2_, there is a clearly visible network in the fluorescence micrographs (Fig. 3A, bottom left) that evolves into a dense network at 3 mol% PtdIns[4,5]P_2_ (Fig. 3A, bottom right) concomitant with an increased skeleton network density at 1 and 2 mol% PtdIns[4,5]P_2_. In addition, the relative bundling of the filaments becomes more prominent (Fig. 3E). It is further evident that an increase in receptor lipid concentration from 1 to 2 mol% PtdIns[4,5]P_2_ increases the number of local domains with nematic order (Fig. 3A, middle and left). The calculated average nematic order obtained for *n* fluorescence micrographs and *m* preparations clearly indicate that in the absence of POPS no significant nematic ordering occurs (Fig. 3B). In contrast, *q*_mean_ calculated for networks on SLBs doped with POPS increased from 0.26 *±* 0.02 (median *±* median absolute deviation) at 1 mol% to 0.36 *±* 0.07 at 3 mol% PtdIns[4,5]P_2_, whereas *q*_mean_ remains between 0.24 *±* 0.02 and 0.27 *±* 0.01 in the absence of POPS.

**Fig. 3.**
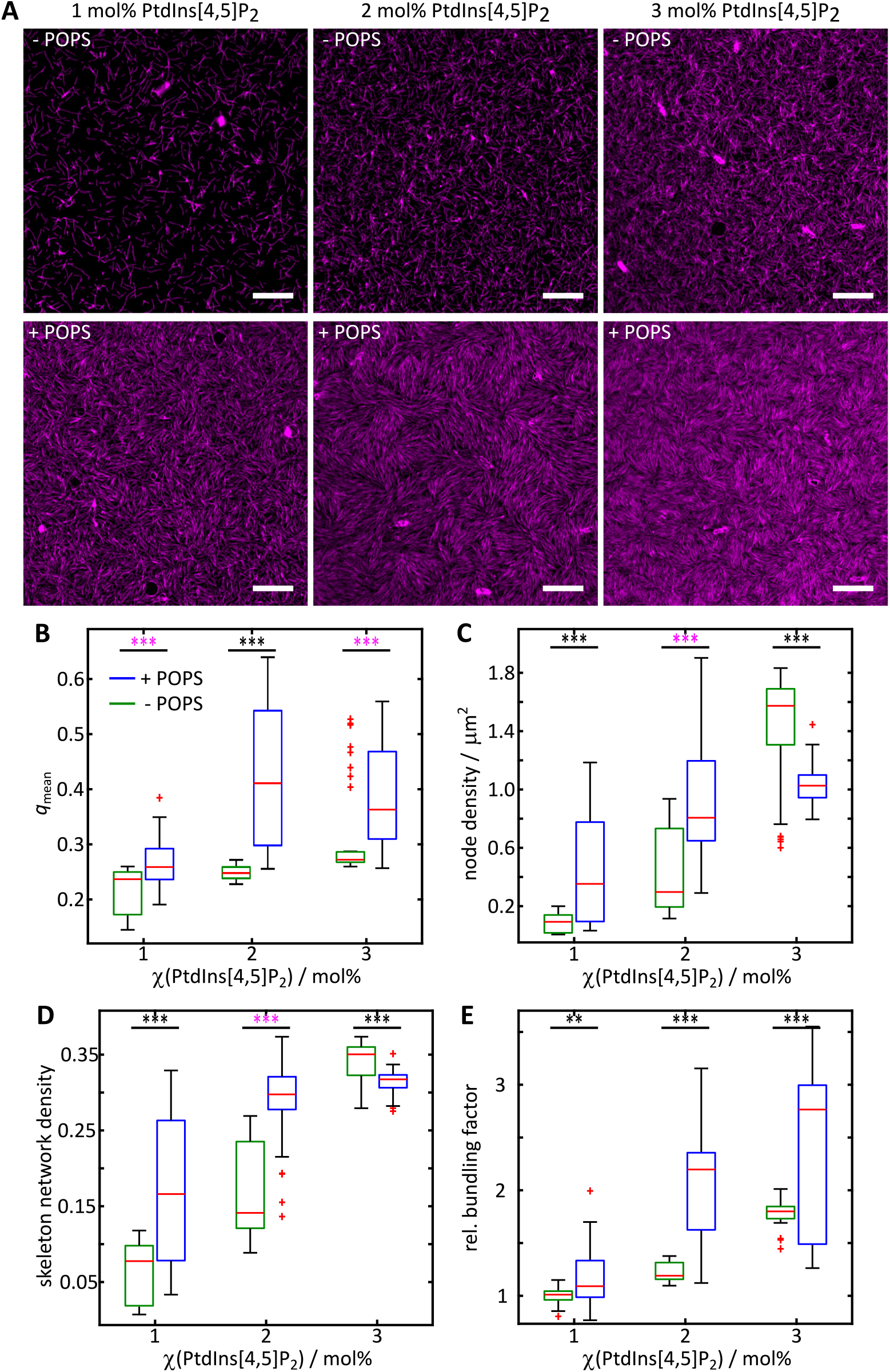
Comparison of the F-actin network self-organization of MACs as a function of PtdIns[4,5]P_2_ and POPS content. (**A**) Exemplary fluorescent micrographs for the F-actin (magenta) self-organization on SLBs without POPS (upper row) and doped with 17 mol% POPS (lower row) and varying PtdIns[4,5]P_2_ concentrations. Scale bars: 10 µm. (**B**) Mean nematic order parameter (*q*_mean_), (**C**) node density, (**D**) skeleton network density and (**E**) relative bundling factor of F-actin networks bound to SLBs without POPS (green) and doped with 17 mol% POPS (blue) as a function of the PtdIns[4,5]P_2_ content. For the analysis, *n* images of *m* independent preparations were evaluated. 1 mol% (*n* = 30, *n*_q,mean_ = 30, *m* = 4), 2 mol% (*n* = 23, *n*_q,mean_ = 24, *m* = 3) and 3 mol% (*n* = 34, *n*_q,mean_ = 39, *m* = 4) PtdIns[4,5]P_2_ ; 1 mol% (*n* = 34, *n*_q,mean_ = 37, *m* = 4), 2 mol% (*n* = 60, *n*_q,mean_ = 52, *m* = 6) and 3 mol% (*n* = 40, *n*_q,mean_ = 56, *m* = 6) PtdIns[4,5]P_2_ + 17 mol% POPS. Boxes ranging from 25^th^ to 75^th^ percentiles of the sample, while whiskers represent the most extreme data points not considered as outliers (red crosses). Medians are shown as red horizontals within the boxes. Statistical *t* -test: **: *p ≤* 0.01, ***: *p ≤* 0.001; *Welch*-test: ***: *p ≤* 0.001.

Another parameter that characterizes the actin networks is the node density (Fig. 3C). In the absence of POPS, the projected node density (Fig. 3C) shows the same trend as the skeleton network density (Fig. 3D) as expected for an isotropic network on the membrane surface (Fig. 3A, top row). The node density increases from 0.09 *±* 0.06 µm^*−*^ _1_ (1 mol% PtdIns[4,5]P_2_) to 1.57 *±* 0.18 µm^*−*^ _1_ (3 mol% PtdIns[4,5]P_2_). However, in the presence of POPS, the node density is initially larger than in the absence of POPS (4-fold increase at 1 mol% PtdIns[4,5]P_2_ and 3-fold increase at 2 mol%), but already at 3 mol% PtdIns[4,5]P_2_ the density reduces due to the parallel orientation of the filaments, which leads to the formation of domains with nematic order.

### Myosin II induced reorganization of membrane bound minimal actin cortices

In the cortex of living cells, connectivity of the actin network and myosin motor activity go hand in hand constituting a pre-stressed and contractile network capable of performing various tasks that require quick shape changes of the cell such as adhesion, migration or division. Here, we investigated how myosin motors reorganize membrane-bound actin filaments as a function of PtdIns[4,5]P_2_ and POPS content. Bipolar myosin II filaments were added to a pre-assembled F-actin network attached via ezrin T567D on PtdIns[4,5]P_2_ doped SLBs (Fig. 4A). Dual color fluorescence microscopy images were acquired using total internal reflection fluorescence (TIRF) microscopy, localizing both actin filaments (magenta) and myosin motors (green). The addition of pre-polymerized myosin II filaments and ATP to the MACs on SLBs without POPS (Fig. 4B, -10 s) led to fast myosin II binding within a few seconds (Fig. 4B, +20 s). Upon binding, myosin II clusters form leading only to minor network reorganization (Fig. 4B, +60-600 s), i.e., large scale re-organization of F-actin such as global or local contraction of the network does not occur (Movie S1). However, if the MAC is bound onto POPS-doped membranes, a fast F-actin reorganization is observed (Fig. 4B, +20 s) upon myosin II addition (Fig. 4B, -10 s) leading to the occurrence of local actin filament clusters with dense myosin centers, so-called asters (Fig. 4B, +60-600 s). Rapid restructuring of the actin network into asters is accompanied by a loss of F-actin fluorescence intensity in their vicinity, leading eventually to local network contraction (Movie S2).

**Fig. 4.**
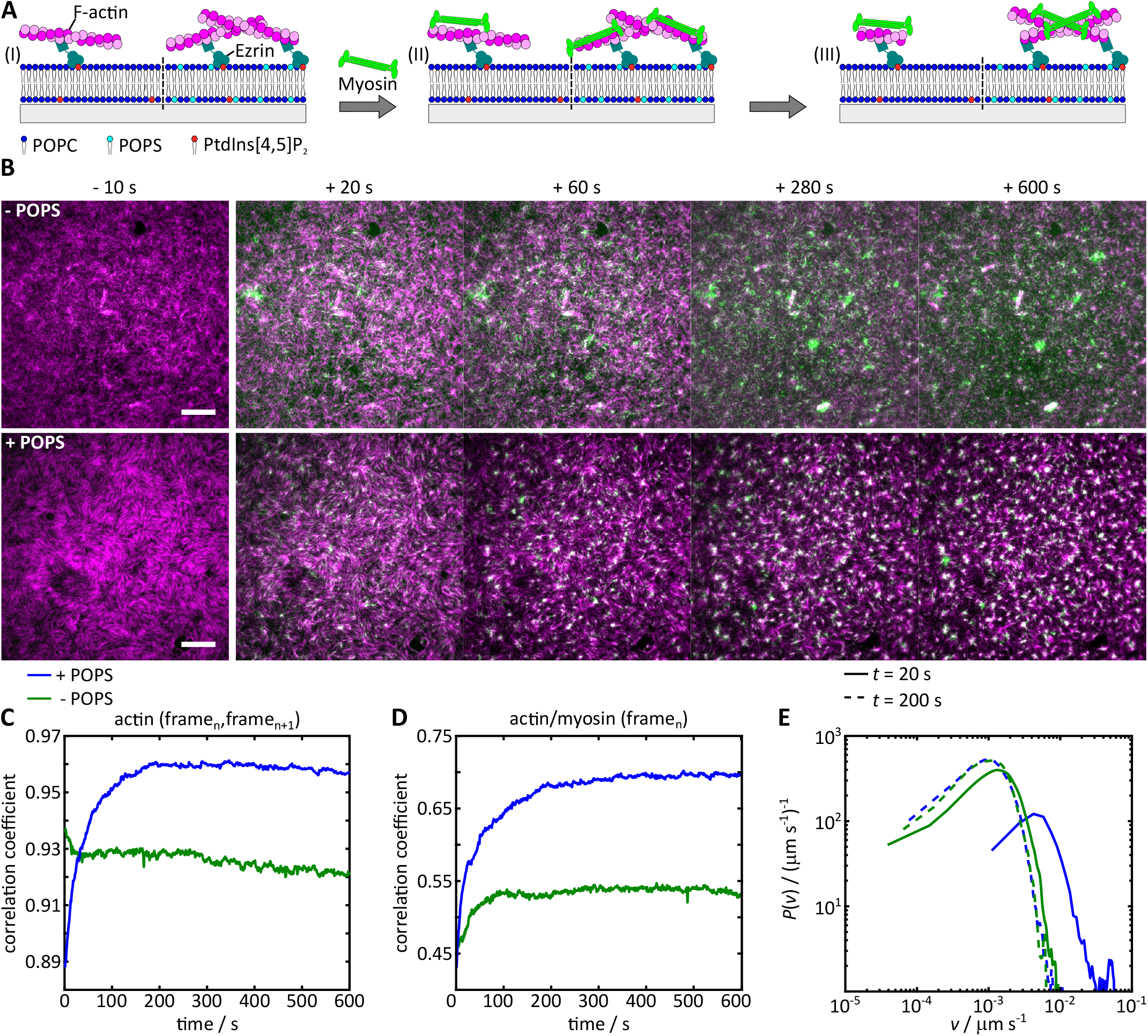
Myosin II induced reorganization of membrane bound minimal actin cortices. (**A**) Schematic illustration of myosin II induced reorganization of F-actin networks bound to SLBs composed of POPC, PtdIns[4,5]P_2_ (left) and POPS (right). (I) Membrane bound MAC prior to myosin II addition, (II) immediately after myosin II binding and (III) after reorganization. (**B**) Exemplary fluorescent micrographs of F-actin networks (magenta) bound to SLBs (*χ*(PtdIns[4,5]P_2_ = 3 mol%) without POPS (top row) and doped with 17 mol% POPS (bottom row) prior (−10 s) and after (+20-600 s) myosin II (light green) addition. Scale bars: 10 µm. Time profiles of the samples shown in (B) for the (**C**) 2D cross-correlation coefficient of the F-actin fluorescence intensity from each frame (frame) to the following frame (frame_n+1_) and (**D**) for the F-actin fluorescence intensity to myosin II fluorescence intensity within the same frame. (**E**) Velocity magnitude distribution of the samples shown in (B) 20 s (solid line, active state) and 200 s (dashed line, passive state) after myosin binding.

To quantify myosin II binding over time, we monitored the time-dependent normalized fluorescence intensity of myosin II (Fig. S2B). Both time traces, in the absence and presence of POPS, saturate after 100 s, indicating that myosin II binding is nearly completed. Simultaneously, the fluorescence intensity of F-actin decreases (Fig. S2A), which can be attributed to bundling, compaction and removal of the filaments from the membrane surface (22). It occurs in both cases, in the presence and absence of POPS and independent of whether the network contracts or not.

Image cross-correlation was employed to assess the emergence of a dynamic steady state and the extent of F-actin reorganization due to myosin activity and to quantify the differences between a non-contracting and a locally contracting MAC. Therefore, the actin fluorescence signal in each frame (frame_n_) was correlated with the actin fluorescence signal in the following frame (frame_n+1_) (Fig. 4C). For the non-contracting F-actin network bound to a POPS-free membrane, the correlation coefficient drops only slightly in the first few seconds as expected for the initial disassembly of the F-actin network upon myosin binding before stabilizing at about 0.93, indicating that no large scale reorganization of F-actin into asters occurs, i.e., a dynamically stable prestressed network results. In contrast, the correlation coefficient for the contracting network in the presence of POPS increases from 0.89 to about 0.96 over 200 s. Within the same time period, the cross-correlation of the F-actin and myosin II fluorescence signal in the same frame (frame_n_) (Fig. 4D) increases from 0.45 to about 0.68 indicative of a contractile network. In addition to the two extremes of a contracting and non-contracting network, we also found cases in which aster formation occurs only very locally, while other parts show minor reorganization of the network. We refer to these re-organization processes to a partially contracting network (Movie S3).

Contractile activity of the actomyosin network is further assessed by determining the velocity magnitude distribution obtained from particle image velocimetry (PIV) (Fig. 4E). Comparison of the velocity probability distributions 20 s (solid lines) and 200 s (dashed lines) after binding of myosin II for the experiments shown in Fig. 4B demonstrates that contractile networks have higher velocities at the onset of reorganization (Fig. 4E, blue solid line), while non-contractile systems remain in the lower velocity regime (Fig. 4E, green solid line), similar to the situation at 200 s.

Based on the criteria of contracting and non-contracting MACs as shown in Fig. 4, we classified for each preparation, whether the network contracts, partially contracts or does not contract (Fig. S3). Fig. 5 shows the fraction of membrane bound MACs belonging to the three different classifications as a function of POPS clearly indicating that the presence of POPS shifts the system in the direction of local contraction. In the absence of POPS, partial or full contraction can only be induced at much larger PtdIns[4,5]P_2_ concentrations of 5 or 8 mol% (Fig. S4). Apparently, POPS contributes significantly to the contractility of the actin network. Two main aspects are presumably responsible for this behavior as discussed in the next section.

**Fig. 5.**
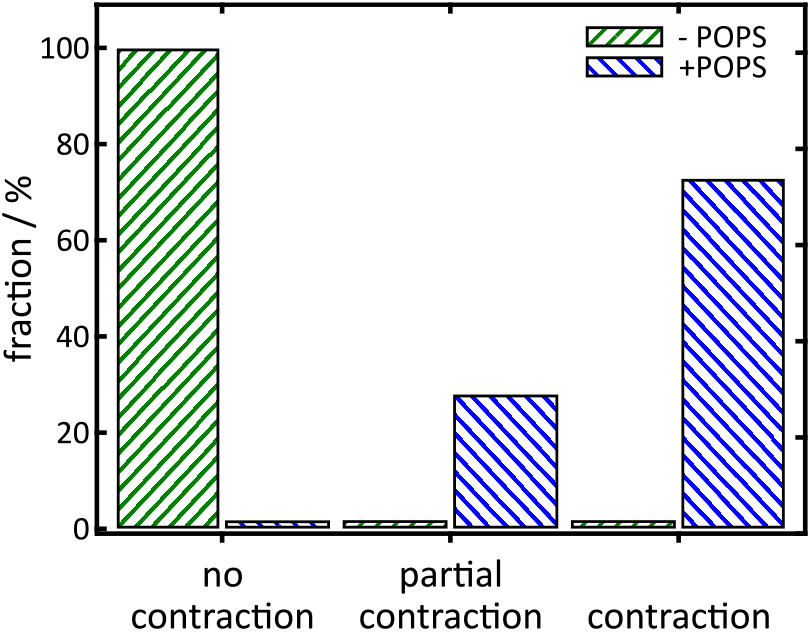
Dependence of the lipid membrane composition on the myosin II induced MAC contractility. Fraction of membrane bound MACs with a receptor lipid content of 1-3 mol% PtdIns[4,5]P_2_ showing no, partial or complete network contraction upon myosin II addition as function of the POPS content. For the analysis, *n* contraction experiments were evaluated. No contraction: 2 mol% (*n* = 5) and 3 mol% (*n* = 8) PtdIns[4,5]P_2_. Partial contraction: 3 mol% (*n* = 3) PtdIns[4,5]P_2_ + 17 mol% POPS. Complete contraction: 1 mol% (*n* = 2), 2 mol% (*n* = 5) and 3 mol% (*n* = 1) PtdIns[4,5]P_2_ + 17 mol% POPS.

## Discussion

The emergence of cortical tension is a hallmark of eukaryotic cells enabling them to maintain their shape, divide, migrate and to rapidly respond to external cues (34). The stresses in the shell arise due to the presence of myosin motors within a dense, reversibly cross-linked actin cytoskeleton. Local changes in cortex composition and architecture can lead to gradients, which may result in local contractions (2). In contractile gels, Koenderink and coworkers (35) identified a narrow regime of motor activity and cross-linker density in which the F-actin network displays contractile behavior. The role of the plasma membrane in this process is, however, largely unexplored. It not only acts as a dielectric barrier for the exchange with the environment but also provides a cue to regulate contractility of the cortex keeping contractility in a regime that permits the cell to allow the transition from a pre-stressed homeostatic state to cortical flow induced by local contractions but also to avoid a global collapse of the network. Murrell and coworkers simply crowded an F-actin network onto a SLB by methyl-cellulose (19, 20) and found, by correlating network and filament properties, that the entropy production rate of actomyosin networks is highest in non-contractile, stable networks (31). Vogel et al. (21) artificially attached F-actin via biotin-neutravidin to the SLB. Upon F-actin contraction, driven by myosin II motors, the filaments fragmented i.e, buckling and breaking was observed. With a histidine-tagged VCA-domain bound to Ni-NTA-DGS, Ganziger et al. (22) could induce Arp2/3 branched actin polymerization onto the membrane. These actomyosin cortices were characterized by a dynamic steady state process of myosin II induced fragmentation/contraction and actin polymerization. An engineered membrane–actin linker constructed from the actin-binding domain of ezrin attached by Ni-NTA-DGS was used by Köster and coworkers (11, 23, 24). They observed distinct states of actomyosin organization at the membrane surface upon complete ATP consumption. However, even though a few seminal studies have been performed to address the presence of a lipid bilayer, the natural lipid composition and linkage of the F-actin network to the membrane has been neglected. The inner leaflet of the plasma membrane contains phosphatidylserine (PS) being responsible for a fairly large surface charge density (25). While the function of PS, e.g. for homeostasis and as a hallmark of apoptosis, has been extensively studied, its importance for cortex structure and mechanics remains largely unexplored.

By using SLBs we were able to mimic the composition of the inner plasma membrane to shed light not only on the influence of the PtdIns[4,5]P_2_ concentration on membranebound F-actin and actomyosin networks, but also on the impact of PS. Linkage of the F-actin network to the SLBs was realized using the phospho-mimetic mutant of ezrin (ezrin T567D) specifically binding to PtdIns[4,5]P_2_ (Fig. 1C, Fig. S1). With this system in hand, we found that not only the number of reversible contacts provided by ezrin binding to PtdIns[4,5]P_2_ matters but also the lipid composition itself. We observed that in the presence of PS, ezrin binding is more efficient, especially in the physiologically relevant regime of 1-3 mol%. The change in optical thickness is substantially increased in the presence of PS compared to a more neutral membrane (Fig. 1C), where only PtdIns[4,5]P_2_ contributes to the negative surface charge density. We cannot rule out that the number of PtdIns[4,5]P_2_ binding sites is reduced in the absence of POPS (36, 37), but we mainly assume that the increased protein thickness in presence of POPS is due to a higher packing density of ezrin T567D and an altered protein orientation fostered by a conformational constraint provided by the presence of negative charges in the bilayer. At 1-3 mol% PtdIns[4,5]P_2_ in pure POPC membranes, the T567D ezrin mutant is known to bind in a less packed conformation to the membrane (8). At larger PtdIns[4,5]P_2_ concentrations, a higher packing density is reached. In this regime (*χ*(PtdIns[4,5]P_2_) *≥* 6 mol%) and in the absence of POPS, the ΔOT = *nd* value translates into a physical protein layer thickness of *d* = 3.2 *±* 0.4 nm (mean *±* standard deviation) assuming a refractive index of *n* = 1.455 (38)) being in good agreement with the height of a densely packed membrane-bound ezrin layer (3.0 *±* 0.4 nm) as determined by atomic force micrographs (8). In the presence of POPS (*χ*(PtdIns[4,5]P_2_) *≥* 6 mol%), *d* reads 4.1 *±* 0.4 nm being about 30 % larger suggesting a different organization of ezrin T567D on the membrane. It is conceivable that the different organization is a result of repulsive interactions between the negatively charged membrane surface and the negative net charge of the protein (4) at pH 7.4 (https://www.protpi.ch/).

Concomitantly with the mode of ezrin attachment, the actin density and architecture is different in the 1-3 mol% PtdIns[4,5]P_2_ regime (Fig. 3A). Not only the F-actin density is slightly higher in the presence of POPS as indicated by the node density (Fig. 3C) (28), the skeleton network density (Fig. 3D) and the relative bundling factor (Fig. 3E), but also the order of the F-actin networks differs (Fig. 3B). We observed a tendency of the network to form nematic structures at 1-3 mol% PtdIns[4,5]P_2_ and in the presence of POPS. We attribute the emergence of nematic phases to repulsion between the negatively charged membrane surface and the actin filaments, which allows the F-actin to pass over bundles into the nematic phase. According to Onsager (39) and the extended model of Khokholov and Semenov (40) for semiflexible rods like F-actin, the formation of nematically aligned filaments depends on the critical packing density at which filaments align and thereby reduce their excluded volume (41). While the critical F-actin concentration in solution can range between 75-100 µM (42), nematic phases can also be formed at low concentrations (2.3-5 µM) (31, 43) if crowding agents are applied. As we observed the transition from an isotropically to a nematically organized network at an actin concentration of only about 1.7 µM, and without any crowding agent, we conclude that binding of F-actin in a quasi-twodimensional network on the membrane can readily lead to a local packing density, which is above the critical concentration needed for filament alignment. In the absence of POPS, a larger PtdIns[4,5]P_2_ concentration is required to induce this transition (Fig. S5A-C).

More importantly, only in the presence of POPS at 1-3 mol% PtdIns[4,5]P_2_, we find a contractile network with local aster formation (Fig. 5) and an increased contraction velocity (Fig. 4E) when we add myosin II motors. However, if the PtdIns[4,5]P_2_ concentration is increased up to 8 mol% in the absence of POPS to achieve a higher membrane charge density that matches the charge provided by PS and a greater network density or order (Figs. S4 and S5C-E), contraction is still largely impaired. A similar observation was made by Murrell and Gardel (19). They found that the attachment of a contractile actomyosin network to a bilayer containing > 8 mol% of FimA2 anchors attached via a His-tag compromises the ability of the network to form contraction foci, in contrast to a network that is only crowded onto the membrane with methyl-cellulose.

In the presence of POPS and only moderate anchoring via 1-3 mol% PtdIns[4,5]P_2_, the network may slide on the surface and can form contraction foci with less resistance, in part given by the repulsion between the membrane and the actin filaments (44, 45). Simply increasing the anchoring points to the membrane by increasing the PtdIns[4,5]P_2_ concentration and thus the ezrin coverage appears not to be sufficient to create local contraction. Connectivity to the membrane is too large and higher motor activity would be required to overcome the constraints. The increased friction of the lipid PtdIns[4,5]P_2_ in the bilayer requires larger motor forces to overcome the drag necessary for contraction. However, the formation of nematic order does not impair local contraction. Contraction in non-sarcomeric structures is especially large in the nematic phase with dense two-dimensional F-actin bundles with apolar orientation (19). Blanchoin and coworkers (46) showed that antiparallel bundles of F-actin display substantially faster contraction compared to non-ordered actin meshes using in vitro contractility assays. To examine this aspect in our system, we plotted *q*_mean_ as a function of contractility (Fig. S6) and indeed found a correlation between contractility and nematic domains. Hence, in the context of local contractility of actin cortices, the natural occurrence of PS may fulfill the task of a repulsive boundary that enables the network to hover/lift above the bilayer. Thereby both higher network densities and higher order of F-actin is achieved enabling contractility in otherwise too diluted networks. Indeed, in the absence of POPS at low but physiological PtdIns[4,5]P_2_ concentrations (1-3 mol%) asters were not found. Not only less ezrin is bound to the membrane leading to a lower amount of F-actin on the bilayer but also less motility and compliance (the area compressibility modulus rises due to suppressed fluctuations) due to missing repulsion is accomplished. As a consequence, contractility is impaired and proper biological function compromised. In conclusion, the lipid composition is a crucial variable often overlooked when studying actomyosin structure and contractility of the cortex. Our results illustrate that lipids which are not involved in the direct binding of cytoskeletal structures greatly influence connectivity and contractility of F-actin networks. Given the concentrations of PtdIns[4,5]P_2_ (1 mol%) (32) and POPS (about 17 mol%) (26, 33) in the plasma membrane, they appear to be ideally chosen to meet the requirements for connectivity and contractility.

## Methods and Materials

### Vesicle preparation

1-10 mg/ml stock solutions of 1-palmitoyl-2-oleoyl-*sn*-glycero-3-phosphocholine (POPC), 1-palmitoyl-2-oleoyl-*sn*-glycero-3-phospho-L-serine (POPS, Avanti Polar Lipids, Alabaster, AL, USA) and ATTO 390-1,2-dipalmitoyl-*sn*-glycero-3-phosphoethanolamine (Atto 390-DPPE, ATTO-TEC, Siegen, Germany) were prepared in chloroform. L-*α*-phosphatidylinositol-4,5-bisphosphate (PtdIns[4,5]P_2_, brain porcine, Avanti Polar Lipids, Alabaster, AL, USA) was freshly dissolved in chloroform/methanol/H_2_O (10:20:8) to a final concentration of 1 mg/mL. Lipid mixtures (0.4 mg) were prepared in chloroform/methanol (10:1) and organic solvents were evaporated with a nitrogen stream followed by 3 h in vacuum. The dried lipid films were stored at 4 °C until needed.

Small unilamellar vesicles (SUVs) were prepared by re-hydrating a lipid film in spreading buffer (50 mM KCl, 20 mM Na-citrate, 0.1 mM NaN_3_, 0.1 mM ethylenediaminetetraacetic acid (EDTA), pH 4.8) (47), incubating for 30 min, subsequent vortexing 3 × 30 s at 5 min intervals and a final sonification step for 30 min at room temperature (cycle 4 60 %, Sonopuls HD2070, resonator cup; Bandelin, Berlin, Germany). PtdIns[4,5]P_2_ containing SUVs were used immediately for preparation of SLBs to avoid PtdIns[4,5]P_2_ degradation (48),

### SLB preparation

Solid supported lipid bilayers (SLBs) were prepared on glass substrates (no. 1.5, Marienfeld-Superio, Lauda-Königshofen, Germany), used for the preparation of MACs, and on silicon wafers coated with 5 µm SiO_2_ (Silicon Materials, Kaufering, Germany), used for reflectometric interference spectroscopy (RIfs). Both substrates were treated for 20 min with a H_2_O/NH_3_/H_2_O_2_ (5:1:1, v/v) solution at 70 °C and subsequently activated for 30 s with O_2_-plasma (Zepto LF PC, Diener electronic, Ebhausen, Germany). The hydrophilized substrates were mounted in a measuring chamber and immediately incubated with SUVs. For MACs on glass slides, SLBs were formed by incubating the substrates for 1 h with SUVs (*m* = 0.2 mg, *c* = 0.53 mg/mL) at 20 °C and excess lipid material was removed by a 10-fold buffer exchange with spreading buffer followed by ezrin buffer (50 mM KCl, 20 mM Tris, 0.1 mM NaN_3_, 0.1 mM EDTA, pH 7.4). For SLB formation on silicon substrates, SUVs (*m* = 0.2 mg, *c* = 0.53 mg/mL) were spread while the optical thickness was read out. After successful SLB formation, excess lipid material was removed by rinsing 5 min with spreading buffer and 5 min with ezrin buffer.

### Ezrin binding monitored by RIfS

Reflectometric interference spectroscopy (RIfS) is a noninvasive label-free technique to determine optical layer thicknesses (OT = *n d*). The experimental setup was described previously (49). Ezrin T567D was recombinantly expressed in *E. coli* (BL21(DE3)pLysS, Novagen, Madison, WI, USA) and purified as described previously (29). SLB formation and ezrin binding were monitored using a Flame-S-UV/vis spectrometer (Ocean Optics, Dunedin, FL, USA), recording a spectra every 2 s and analyzed utilizing a custom MATLAB script (R2021a, Mathworks). After SLB formation, the surface was passivated with a BSA solution (1 mg/mL in ezrin buffer) for 5 min. After rinsing the system for another 5 min with ezrin buffer, ezrin T567D was added (0.8 µM) for 10 min. Unbound protein was removed by rinsing with ezrin buffer.

### Preparation of MACs

Ezrin T567D was bound to the SLBs at a concentration of 1 µM overnight at 4 °C. Excess protein was removed by a 10-fold buffer exchange with ezrin buffer and F-actin buffer (50 mM KCl, 20 mM Tris, 2 mM MgCl_2_, 0.1 mM NaN_3_, pH 7.4). For F-actin pre-polymerization ATTO 594-NHS ester (ATTO-TEC, Siegen, Germany) labeled non-muscle G-actin and unlabeled monomers (Cytoskeleton, Denver, CO, USA) were solved in a 1:10 ratio and a final concentration of 0.44 mg/mL in G-buffer (5 mM Tris, 0.2 mM CaCl_2_, 0.1 mM NaN_3_, pH 8.0). Actin oligomers were depolymerized by the addition of dithiothreitol (DTT, 0.5 mM) and adenosine 5^*′*^-triphosphate (ATP, 0.2 mM) for 1 h on ice. Remaining actin aggregates were centrifuged (17000 × *g*, 20 min, 4 °C) and polymerization was induced by the addition of 10% of the total volume of polymerization solution (500 mM KCl, 20 mM MgCl_2_, 20 mM ATP, 50 mM guanidine carbonate, pH 7.4). After a polymerization time of 20 min at 20 °C, the F-actin solution was mixed with unlabeled phalloidin in a 1.5 % (*n*/*n*) ratio and incubated for another 20 min. MACs were formed at 20 °C by incubating the ezrin T567D decorated SLBs with polymerized F-actin at a concentration of 4.6 µM for at least 2 h. Unbound filaments were washed off by a 10-fold buffer exchange with F-actin buffer.

### Contraction experiments

Myosin II was purified from rabbit skeletal muscle and fluorescently labeled with Dye-Light 488 (Invitrogen, Carlsbad, CA, USA) according to (50). Labeled and unlabeled myosin II were stored separately in myosin storage buffer (300 mM KCl, 25 mM KH_2_PO_4_, 0.5 mM DTT, 50 % (*v*/*v*) glycerol, pH 6.5), where the high ionic strength prevents myosin self-assembly into bipolar filaments. For experiments, myosin II was dialyzed overnight in glycerol free myosin buffer (300 mM KCl, 20 mM imidazol, 4 mM MgCl_2_, 1 mM DTT, pH 7.4) and controlled self-assembly into bipolar filaments was induced by adjusting a KCl concentration of 50 mM via mixing with myosin polymerization buffer (20 mM imidazol, 1.6 mM MgCl_2_, 1 mM DTT, 1.2 mM Trolox, pH 7.4). After an incubation time of 10 min at 20 °C, the bipolar myosin II filaments were immediately used for contractile experiments.

For the contraction experiments, MACs were transferred into an actomyosin buffer by a 10-fold buffer exchange (50 mM KCl, 20 mM imidazol, 2 mM MgCl_2_, 1 mM DTT, 1 mM Trolox, pH 7.4). The reorganization of MACs was performed at a final ATP concentration of 0.1 mM combined with a ATP-regeneration system of creatine phosphate (10 mM)/creatine kinase (0.1 mg/mL) (50) and a myosin II concentration of 0.4 µM.

### Image acquisition

Confocal fluorescence images were acquired with the upright confocal laser scanning microscope LSM 880 (CLSM, Carl Zeiss Microscopy GmbH, Oberkochen, Germany) using a 40× objective (W Plan-Apochromat M27, NA = 1.0, Carl Zeiss Microscopy GmbH, Oberkochen, Germany). The membrane dye ATT0 390-DPPE was excited at *λ*_*ex*_ = 405 nm (diode laser, 30 mW) and detected at 450 -550 nm. The ATTO 594-labeled F-actin was excited at *λ*_*ex*_ = 561 nm (diode laser, 20 mW) and fluorescence was detected between 600-700 nm, using an Airyscan detector.

Dual color TIRF microscopy was performed using the IXpolre TIRF system (cellTIRF-4Line, Olympus Deutschland GmbH, Hamburg, Germany) equipped with a 100× oil objective (UPLAPO-HR, NA = 1.5, Olympus Deutschland GmbH, Hamburg, Germany). The ATTO 594-labeled F-actin was excited at *λ*_*ex*_ = 561 nm (diode laser, 100 mW) and DyLight 488-labeld myosin II at *λ*_*ex*_ = 488/491 nm (diode laser, 500/100 mW). Exposure times were 20 ms (myosin II) and 40 ms (actin), fluorescence signals were detected using a Zyla 4.2 sCMOS (Andor Technology Ltd., Belfast, UK).

### Image analysis

Fluorescence micrographs of membrane-bound MACs were skeletonized by means of the Jupyter notebook (51) based custom written “*tube filter*” analysis. In the first step of the analysis, a contrast limited adaptive histogram equalization (CLAHE) (52) and a two dimensional Gaussian-filter were used to equilibrate the global image intensity, improve the local image contrast and reduce high-frequency noise. Based on the F-actin intensity, this two-dimensional exposure-corrected micrographs (*I*(*x, y*)) can be described as three-dimensional surfaces. In the second analysis step, the Hessian image matrix (*H*_*I*_ (*x, y*)), eq. 1) of the exposure-corrected micrographs was calculated, containing the second partial derivatives of the input image.

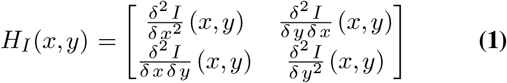

The signal-to-noise enhanced tube-filtered images (Fig. 2B1) were generated by means of the second eigenvalues *λ*_2_(*x, y*) of *H*_*I*_ (*x, y*), describing the minimal surface curvature at each image pixel of *I*(*x, y*). Tube-filtered images were subsequently suited for “conventional” adaptive thresholding and skeletonization.

From the skeletonized images (Fig. 2B2), the skeleton network density, defined as the ratio of filamentous pixels (Fig. 2B2, white) to all images pixels and filament intersections, referred to as node density (Fig. 2B3, red crosses), were determined. A node was defined as a filamentous pixel with more then two adjacent filamentous pixels.

The relative bundling factor calculation was performed via extracting the fluorescence intensity of membrane-bound F-actin, by masking actin fluorescence images with the corresponding skeletonized images (Fig. S7). The actin fluorescence was determined at the overlapping positions, averaged for the respective micrograph and normalized with the mean F-actin intensity at *χ*(PtdIns[4,5]P_2_) = 1 mol% without POPS, assuming only single actin filaments at this conditions.

The local nematic order parameter *q* = 2(cos^2^ *θ −* 1*/*2) of membrane-bound MACs was calculated according to Seara et al. (31), by utilizing the published MATLAB routines. For each fluorescence image the mean nematic order parameter (*q*_mean_) was calculated by averaging all local *q*-values. Since the window size for the generation of the alignment vector field (Fig. 2A2, yellow) significantly influences the calculated *q*-values, *q*_mean_ was computed for each image as a function of the window size in order to determine an optimal one (Fig. S8). The optimal window size used in this work ranged between 1-2 µm.

Actin and myosin II intensities were analyzed after the initial binding of myosin II. The respective sample intensity was averaged for each frame and normalized by means of the maximal intensity within the particular time series. Intensity correlation within one frame (frame_n_) or to the following frame (frame_n+1_) were performed for entire time series utilizing the MATLAB function corr2 (Fig. 4C-D).

For the calculation of the F-actin velocity magnitude distribution of contractile actomyosin networks, the samples were analyzed by means of particle image velocimetry (PIV) using the MATLAB based PIVlab (version 2.53) from Thielicke and Sonntag (53). To compensate for thermal drift, the time series were cropped to the region of interest and subsequently adjusted using the StackReg (http://bigwww.epfl.ch/thevenaz/stackreg/) plugin of ImageJ (https://imagej.nih.gov/ij/). The PIV analysis was performed at a time difference of 10 s between the analyzed frames, using the parameters listed in Table S1. The final F-actin velocity magnitude distribution calculation was performed by averaging the F-actin velocity magnitude per frame and utilizing the MATLAB function ksdensity (Fig. 4E and S3). General data analysis, statistical tests and plotting of graphs was done using MATLAB and custom written scripts. Basic image processing was done with ImageJ. All custom written scripts can be provided on demand.

## Acknowledgments

We like to thank the Koenderink group for the myosin II isolation and protein labeling protocols and J. Gerber-Nolte for technical assistance. The DFG (STE885/18-1 and JA963/19-1, EXC 2067–390729940) and VW foundation (Living Foams) is gratefully acknowledged for financial support. N.L.L. thanks the GGNB for financial support.

