## Supplementary Information for "Impact of native-like lipid membranes on the architecture and contractility of actomyosin networks"

DRAFT

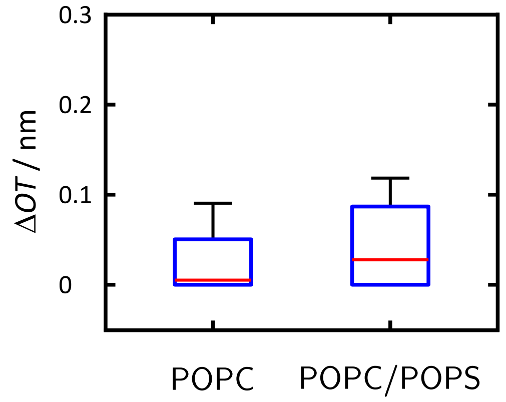


**Fig. 1.** Change in optical thickness ($\Delta OT$) caused by ezrin T567D adsorption on SLBs composed of pure POPC ($m$ = 4) and POPC/POPS (83:17) ($m$ = 4). Negative $\Delta OT$-values caused by the detector noise were set to zero. The median values (red horizontals within the boxes) are 0.02 $\pm$ 0.04 nm (POPC) and 0.04 $\pm$ 0.05 nm (POPC/POPS). Boxes range from 25$^{th}$ to 75$^{th}$ percentiles of the sample, while whiskers represent the most extreme data points.

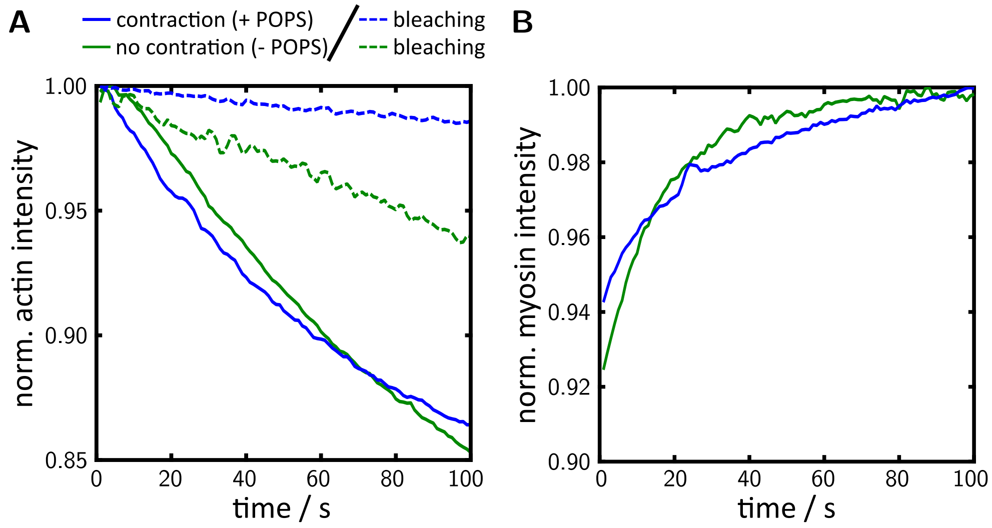


**Fig. 2.** Time-dependent normalized fluorescence intensities of F-actin (**A**, solid lines) and myosin II (**B**) bound to SLBs ($\chi$(PtdIns[4,5]P$_2$ = 3 mol%) without POPS (green, no contraction) and doped with 17 mol% POPS (blue, contraction) after the initial binding of myosin II ($t$ = 0 s). The dashed lines show the general bleaching of the F-actin fluorescence in the absence of myosin (**A**, dashed lines). Time-dependent intensities were extracted from the time series shown in Fig. 4B.

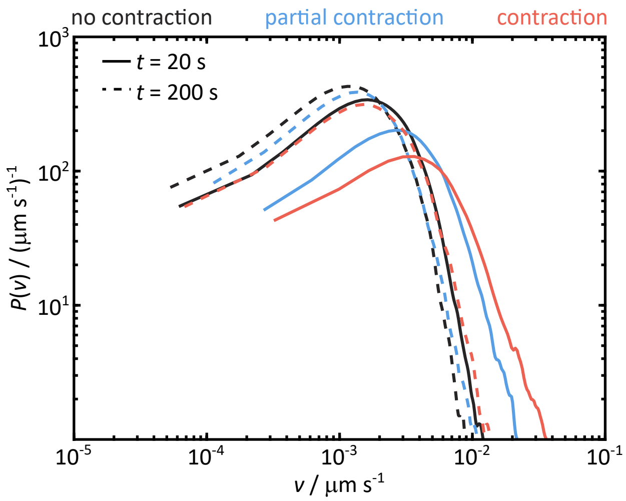


**Fig. 3.** Mean velocity magnitude distribution depending on F-actin network contractility. Comparison of the F-actin network contraction velocity 20 s (solid line, active state) and 200 s (dashed line, passive state), analyzed via PIV, for all non-contracting, partially contracting and fully contracting MACs. For the analysis $n$ time series from individual preparations were used. No contraction ($n$ = 19), partial contraction ($n$ = 7) and full contraction ($n$ = 9).

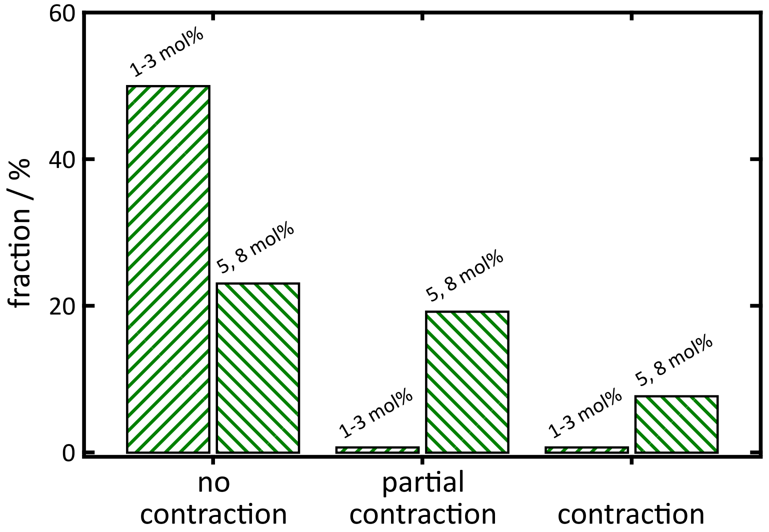


**Fig. 4.** Dependence of the lipid membrane composition on the myosin II induced MAC contractility in the absence of POPS. Fraction of membrane bound MACs with a receptor lipid content of 1-8 mol% PtdIns[4,5]$P_2$ showing no, partial or full network contraction upon myosin II addition. For the analysis, $n$ contraction experiments were evaluated. No contraction: 1-3 mol% ($n$ = 13) and 5, 8 mol% ($n$ = 6) PtdIns[4,5]$P_2$. Partial contraction: 5, 8 mol% ($n$ = 5) PtdIns[4,5]$P_2$. Full contraction: 5, 8 mol% ($n$ = 2) PtdIns[4,5]$P_2$.

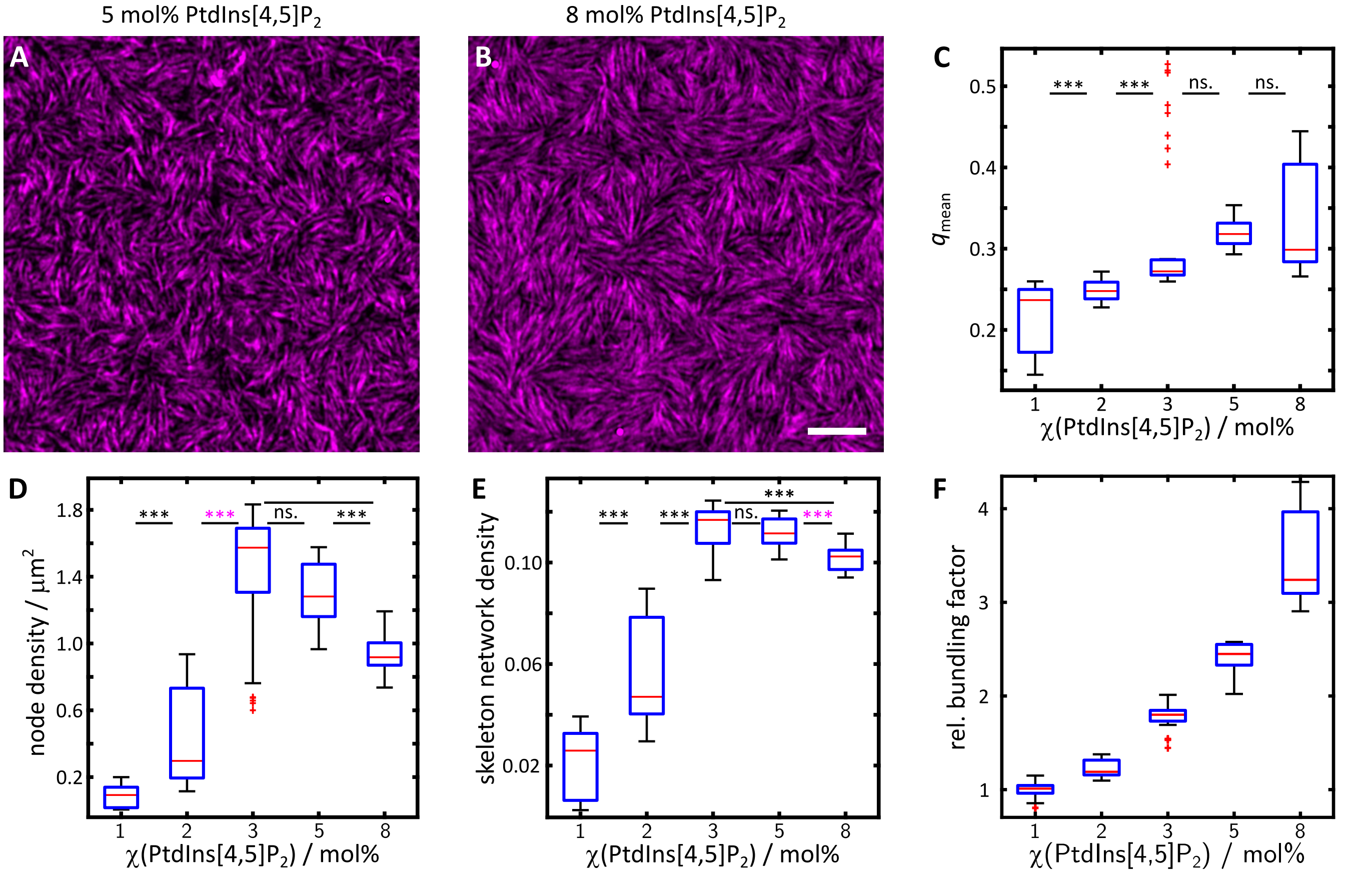


**Fig. 5.** Comparison of the F-actin network self-organization on membranes as a function of PtdIns[4,5]$P_2$ content. Fluorescent micrographs of F-actin (magenta) on SLBs doped with 5 mol% (**A**) and 8 mol% (**B**) PtdIns[4,5]$P_2$ in the absence of POPS. Scale bar: 5 µm. (**C**) Mean nematic order parameter ($q_{\text{mean}}$), (**D**) node density, (**E**) skeleton network density and (**F**) relative bundling factor of F-actin networks bound to SLBs without POPS as a function of PtdIns[4,5]$P_2$ content. For the analysis, $n$ images of $m$ preparations were evaluated. 1 mol% ($n$ = 30, $n_{\text{q,mean}}$ = 30, $m$ = 4), 2 mol% ($n$ = 23, $n_{\text{q,mean}}$ = 24, $m$ = 3), 3 mol% ($n$ = 34, $n_{\text{q,mean}}$ = 39, $m$ = 4), 5 mol% ($n$ = 20, $n_{\text{q,mean}}$ = 20, $m$ = 2) and 8 mol% ($n$ = 31, $n_{\text{q,mean}}$ = 31, $m$ = 3) PtdIns[4,5]$P_2$. Boxes ranging from $25^{\text{th}}$ to $75^{\text{th}}$ percentiles of the sample, while whiskers represent the most extreme data points not considered as outliers (red crosses). Medians are shown as red horizontals within the boxes. Statistical $t$-test: ns.: $p > 0.05$, ***: $p \leq 0.001$; *Welch*-test: ***: $p \leq 0.001$.

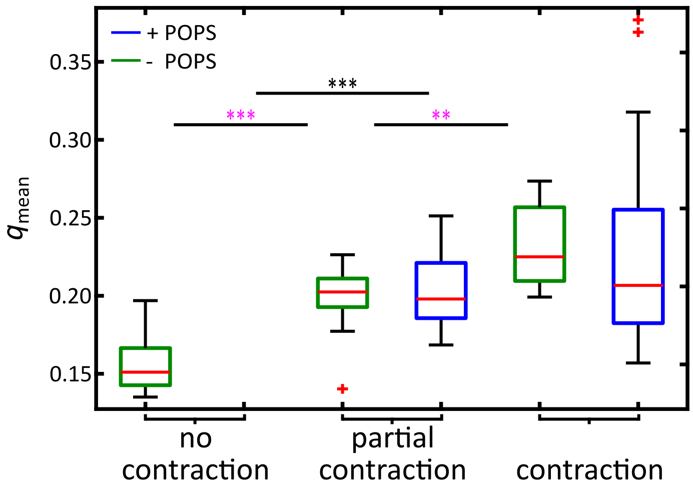


**Fig. 6.** Mean nematic order parameter ($q_{mean}$) of F-actin networks bound to SLBs doped with (blue) and without (green) 17 mol% POPS as a function of the myosin II reorganization. For the analysis *n* images of *m* preparations were used. No contraction ($n$ = 55, $m$ = 19), partial contraction ($n$ = 20, $m$ = 5; $n$ = 12, $m$ = 3) and full contraction ($n$ = 9, $m$ = 2; $n$ = 33, $m$ = 8). Boxes ranging from 25$^{th}$ to 75$^{th}$ percentiles of the sample, while whiskers represents to the most extreme data points not considered outliers (red crosses). Medians are shown as red horizontals within the boxes. Statistical *t*-test: ***: $p \leq 0.001$; *Welch*-test: ***: $p \leq 0.001$, **: $p \leq 0.01$.

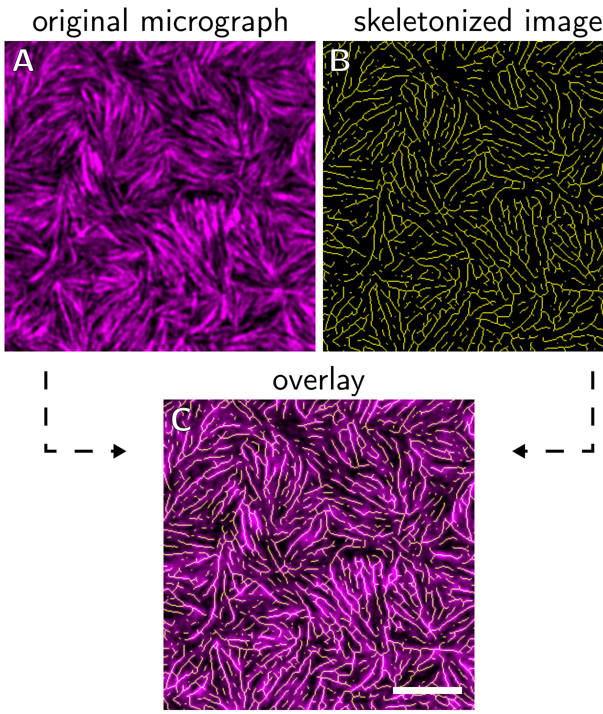


**Fig. 7.** Skeleton based actin intensity and relative bundling factor determination. (**A**) Exemplary fluorescence micrograph of a membrane-bound minimal actin cortex (magenta) and (**B**) of the corresponding skeletonized image (yellow). (**C**) Overlay of the images shown in (A) and (B). Actin fluorescence intensity was read out at the overlapping positions and averaged over the complete micrograph. The relative bundling factor was determined by normalizing the averaged actin intensity with the mean F-actin intensity at $\chi(\text{PtdIns[4,5]P}_2)$ = 1 mol% without POPS.

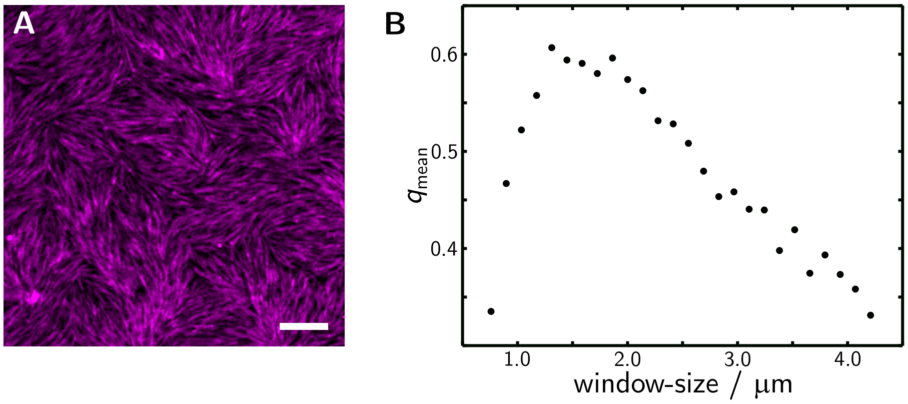


**Fig. 8.** Exemplary determination of the optimal window size for the nematic order parameter. (**A**) Fluorescence micrograph of an exemplary F-actin network (magenta). (**B**) Mean nematic order parameter ($q_{mean}$) of the F-actin network shown in (A) as a function of the applied alignment vector field window size. For the exemplary image in (A) an optimal window size of 1.3 µm ($q_{mean}$ ∼0.6) was determined. Scale bar: 5 µm .

**Table 1.** PIV settings for the calculation of the F-actin velocity magnitude.

| Parameter | |
|---|---|
| CLAHE window-size | 50 px |
| PIV algorithem | FFT window deformation |
| Integration area: | |
| 1. pass | 64 px, step 32 px |
| 2. pass | 32 px, step 16 px |
| 3. pass | 16 px, step 8 px |
| velocity limit | 1 µm $s^{-1}$ |

**Movie 1**: Exemplary time series showing a non contractile minimal actin cortex (magenta) after the addition of myosin II (green, 0 s). Membrane composition: POPC/PtdIns[4,5]$P_2$/ATTO 390-DPPE (96.6:3:0.4)

**Movie 2**: Exemplary time series showing a contractile minimal actin cortex (magenta) after the addition of myosin II (green, 0 s). Membrane composition: POPC/PtdIns[4,5]$P_2$/POPS/ATTO 390-DPPE (79.6:3:17:0.4)

**Movie 3**: Exemplary time series showing a partial contractile minimal actin cortex (magenta) after the addition of myosin II (green, 0 s). Membrane composition: POPC/PtdIns[4,5]$P_2$/POPS/ATTO 390-DPPE (79.6:3:17:0.4)
